# Soil microbiota influences clubroot disease by modulating *Plasmodiophora brassicae* and *Brassica napus* transcriptomes

**DOI:** 10.1101/2020.02.05.935510

**Authors:** Stéphanie Daval, Kévin Gazengel, Arnaud Belcour, Juliette Linglin, Anne-Yvonne Guillerm-Erckelboudt, Alain Sarniguet, Maria J. Manzanares-Dauleux, Lionel Lebreton, Christophe Mougel

## Abstract

The contribution of surrounding plant microbiota to disease development has led to the postulation of the ‘pathobiome’ concept, which represents the interaction between the pathogen, the host-plant, and the associated biotic microbial community, resulting or not in plant disease. The structure, composition and assembly of different plant-associated microbial communities (soil, rhizosphere, leaf, root) are more and more described, both in healthy and infected plants. A major goal is now to shift from descriptive to functional studies of the interaction, in order to gain a mechanistic understanding of how microbes act on plant growth and defense, and/or on pathogen development and pathogenicity. The aim herein is to understand how the soil microbial environment may influence the functions of a pathogen and its pathogenesis, as well as the molecular response of the plant to the infection, with a dual-RNAseq transcriptomics approach. We address this question using *Brassica napus* and *Plasmodiophora brassicae*, the pathogen responsible for clubroot. A time-course experiment was conducted to study interactions between *P. brassicae*, two *B. napus* genotypes, and three soils harboring High (H), Medium (M) or Low (L) microbiota diversities and displaying different levels of richness and diversity. The soil microbial diversity levels had an impact on disease development (symptom levels and pathogen quantity). The *P. brassicae* and *B. napus* transcriptional patterns were modulated by these microbial diversities, and the modulations were dependent of the host genotype plant and the kinetic time. The functional analysis of gene expressions allowed the identification of pathogen and plant-host functions potentially involved in the change of plant disease level, such as pathogenicity-related genes (NUDIX effector) in *P. brassicae* and plant defense-related genes (glucosinolate metabolism) in *B. napus*.

**Author summary:** The untapped soil microbiota diversity can influence plant tolerance and resistance to several pests. A better understanding of the mechanisms underlying the plant / pests / microbiota interaction is required to contribute to the improvement of new plant protection methods taking into account sustainability, respect for the environment, and low input utilization. Our work showed that in the *Plasmodiophora brassicae* / *Brassica napus* pathosystem, the soil microbiota diversity modulated the disease symptom level and the pathogen development. We discovered that soil microbial composition modulated both the pathogen and the plant expression genes profiles. On one hand, the pathogen transcriptome was mainly modulated by the microbial communities at the end of infection, when the pathogen infects a susceptible plant genotype, and the expression of genes potentially involved in growth and pathogenicity was affected. On the other hand, the plant transcriptome was more modulated by the microbial communities at the early step of infection, in the most resistant genotype and the expression of genes potentially involved in defense was affected. This study provides new insights into the molecular basis of soil microbiota-mediated modulation of plant pest diseases.

## Introduction

Plants are constantly interacting with a wide variety of potential pathogens within their environment that can cause serious diseases affecting agriculture. The development of biotic plant diseases depends also on the interaction of both plant and pathogen with the environment. All plant tissues, including leaves [1, 2], seeds [3], and roots [4] are indeed associated with a multitude of microorganisms (viruses, bacteria, archae, fungi, protists, oomycetes, nematodes, protozoa, algae,…) assembled in microbial communities or microbiota. The complex plant-associated microbial community structure and composition, as well as the complex network of interactions between microbial species, are crucial in stress tolerance [5], plant development dynamics [6], yield, nutrition and health [7–10]. This recognition that the plant microbiota may modulate substantially the disease severity and development led to the ‘pathobiome’ postulation, which refers to the pathogenic agent, its surrounding biotic microbial community and their interactions leading to plant disease [11, 12].

In plants, three root-associated microbiota compartments can have a role in the modulation of disease development: the soil microbiota, which represents a great reservoir of biological diversity [13], the rhizosphere corresponding to the narrow zone surrounding and influenced by plant roots [14, 15], and the endosphere (root interior) in which the microbiota diversity is lower than that estimated outside the root [16–19]. Several studies have established close relationships between the rhizosphere microbiome composition and the plant immune system [20–23], the host genotype resistant or susceptible to a pathogen [24], and the life history traits of bioagressors [25], but the mechanisms underlying these relationships have still to be deciphered. It is also known that plants select microbial communities around their roots by specific root exudates [26], that can also function as an additional layer of defense [8]. The defense barrier constituted by recruited microorganisms can be of different types: stimulation of defense-related compounds’ production by the plant, direct antagonism against pathogen (production of antibiotics or antifungal compounds), competition with pathogen for resources [13]. The invasion by a soilborne pathogen led to changes in indigenous plant-associated microbial communities [27, 28] and then in the defense barrier.

Among biotic stress factors, the soilborne plant pathogens cause major yield or quality loss in agricultural crops. This is the case of the protist *Plasmodiophora brassicae,* an obligate biotrophe responsible for clubroot, one of the economically most important diseases of Brassica crops in the world [29]. The life cycle of this soil-borne pathogen can be divided into several phases: survival in soil as spores, root hair infection, and cortical infection [30]. Briefly, during the primary phase of infection, the resting spores germinate in the soil leading to biflagellate primary zoospores that infect the root hairs. In these cells, zoospores multiply to form the primary plasmodia. Secondary zoospores are then released and produce the secondary phase of infection that occurs in the cortex of the roots of the infected plants. During the second phase, multinucleate plasmodia cause the hypertrophy (abnormal cell enlargement) and hyperplasia (uncontrolled cell division) of infected roots into characteristic clubs [31]. These symptoms obstruct nutrient and water transport, stunt the growth of the plant, and consequently reduce crop yield and quality. In root galls, different life cycle stages of *P. brassicae* occur simultaneously.

Transcriptomics studies deciphered in part the mechanisms of the host - *P. brassicae* interaction in simplified experimental conditions, but not in complex soil. During both the spore germination and the primary zoospore stages, the pathogen showed high active metabolisms of chitinous cell wall digestion, starch, citrate cycle, pentose phosphate pathway, pyruvate, trehalose, carbohydrates and lipids [32–34]. During the second phase of infection, genes involved in basal and lipid metabolism were highly expressed [34], as well as the *G-protein-coupled receptors pathway-related* genes [35]. These active metabolic pathways allow *P. brassicae* to take up nutrients from the host cells [30, 36]. During the formation of primary and secondary plasmodia, it is expected that *P. brassicae* secrets an array of effector proteins triggering growth, expansion and differentiation of infected host cells. Nevertheless, few RxLR effectors have been found in *P. brassicae* [32, 37], and no LysM-effectors, known to interfere with chitin detection in fungal-plant interactions [38], were detected. Some candidate potential effectors have however been identified from *P. brassicae* [32, 37, 39], such as Crinkler (CRN) related proteins [40], but their roles in infection and disease development have still to be identified [36]. Only one effector has been characterized in detail: a predicted secreted methyltransferase that can mediate methylation of salicylic, benzoic and anthranilic acids, thereby interfering in the plant salicylic acid-induced defense [41].

Concerning the plant, *P. brassicae* infection altered likewise primary and secondary metabolism, as pathways involved in lipid, carbohydrate, cell wall synthesis, lignification-related genes, arginine and proline metabolism [42–46], producing a sink of plant metabolites assimilated by the pathogen and corresponding to a metabolic cost for the infested plant. Clubroot infection also modified plant hormone homeostasis and defense responses, such as cytokinin biosynthesis, auxin homeostasis, salicylic acid and jasmonic acid metabolism [44–51].

During its life cycle, *P. brassicae* can establish potential relationships with microbiota from soil, rhizospheric soil and roots. Beneficial effect of various specific biocontrol microorganisms in suppressing clubroot has been demonstrated, such as Trichoderma spp. [52], Streptomyces sp. [53, 54], *Heteroconium chaetospira* [55], *Streptomyces platensis* [56], *Bacillus subtilis* [57, 58], *Zhihengliuella aestuarii B18* [59], *Paenibacillus kribbensis* [60], and *Lysobacter antibioticus* [61]. Most of these organisms were isolated from rhizosphere soil or root endosphere. Mechanisms by which these microorganisms protect against clubroot are not yet elucidated but could imply antifungal compounds or molecules up-regulating host plant defense genes. In addition, the microbe abundance in *B. napus* clubroot infected endosphere roots was found higher in asymptomatic roots than in symptomatic roots, and the asymptomatic roots contained many microorganisms with biological control properties and plant growth promotion functions [62]. In Chinese cabbage, invasion by *P. brassicae* modified the rhizosphere and root-associated community assembly during the secondary cortical infection stage of clubroot disease [28]. This shows that the plant microbiota diversity can modulate the plant response to *P. brassicae* and can be considered as a potential reservoir of biocontrol microbe for clubroot prevention. Moreover, in *B. napus*, the plant - microbiota interaction has a role in plant defense against a phytophagous insect (*Delia radicum*) [25, 63].

In order to gain a mechanistic understanding of how soil microbes boost plant growth and defense and/or modulate the pathogen development and pathogenicity, a major challenge is then now to shift from descriptive to functional studies. The aim of this study is to understand how a single root pathogen, *P. brassicae*, interacts with its host, the oilseed rape (*B. napus*), considering the role of the soil microbial diversity as a reservoir of microbial functions related to plant resistance phenotype. To explore how the soil microbial environment may influence the functions of a pathogen and its pathogenesis, and the molecular response of the plant to the infection, we evaluated the effect of different soil microbial diversities obtained by an experimental approach of dilution to extinction on (i) the phenotype of two plant genotypes harboring different levels of susceptibility to the clubroot pathogen, and (ii) the transcriptomes of pathogen and host-plant in interaction.

## Results

### Characterization of the microbial communities in the initial three soil conditions

The microbiological composition after recolonization of the three soils manipulated for having different microbial diversities (High diversity level [H], Medium diversity level [M] or Low diversity level [L]) was analyzed. As expected, the three soils displayed optimal fungal and bacterial densities and similar abundances at the end of recolonization (S1 Fig). Not significant differences for the main soil physicochemical characteristics were observed between the three soils used (S1 Table). The only difference concerned the nitrogen form, that was found mainly in the nitrate form in both H and M and as nitrate and ammonium in L; however, the total nitrogen amount was similar among the three soils, (0.74 to 0.77 g.kg^−1^).

We investigated the effect of the experimental dilution / recolonization on microbiota diversity. Alpha-diversity (within each modality of soil) was analyzed based on the OTUs richness and the Shannon diversity index. For bacterial kingdom (Fig 1A), we observed a statistically significant reduction in richness and specific diversity from H/M to L microbial modalities. For fungal kingdom (Fig 1B), the fungal richness, and to a lesser extent the fungal diversity, decreased also from H to L. Beta-diversity (between soil modalities) was measured for the bacterial and fungal communities (Fig 1C). The soil microbial diversities differed significantly for bacterial and fungal communities. Frequencies of bacterial and fungal phyla, genera and OTUs for each microbial modality are shown in S2 Fig. At the level of phyla, both bacteria and fungi displayed similar frequencies whatever the soil modality, with *Proteobacteria* and *Ascomycota* the dominant phyla, respectively. *Bacillus* and *Pseudomonas* on one hand, and *Schizosaccharomyces* and *Fusarium* on the other hand, were major genera concerning bacteria and fungi, respectively, for the three soils.

**Fig 1.**
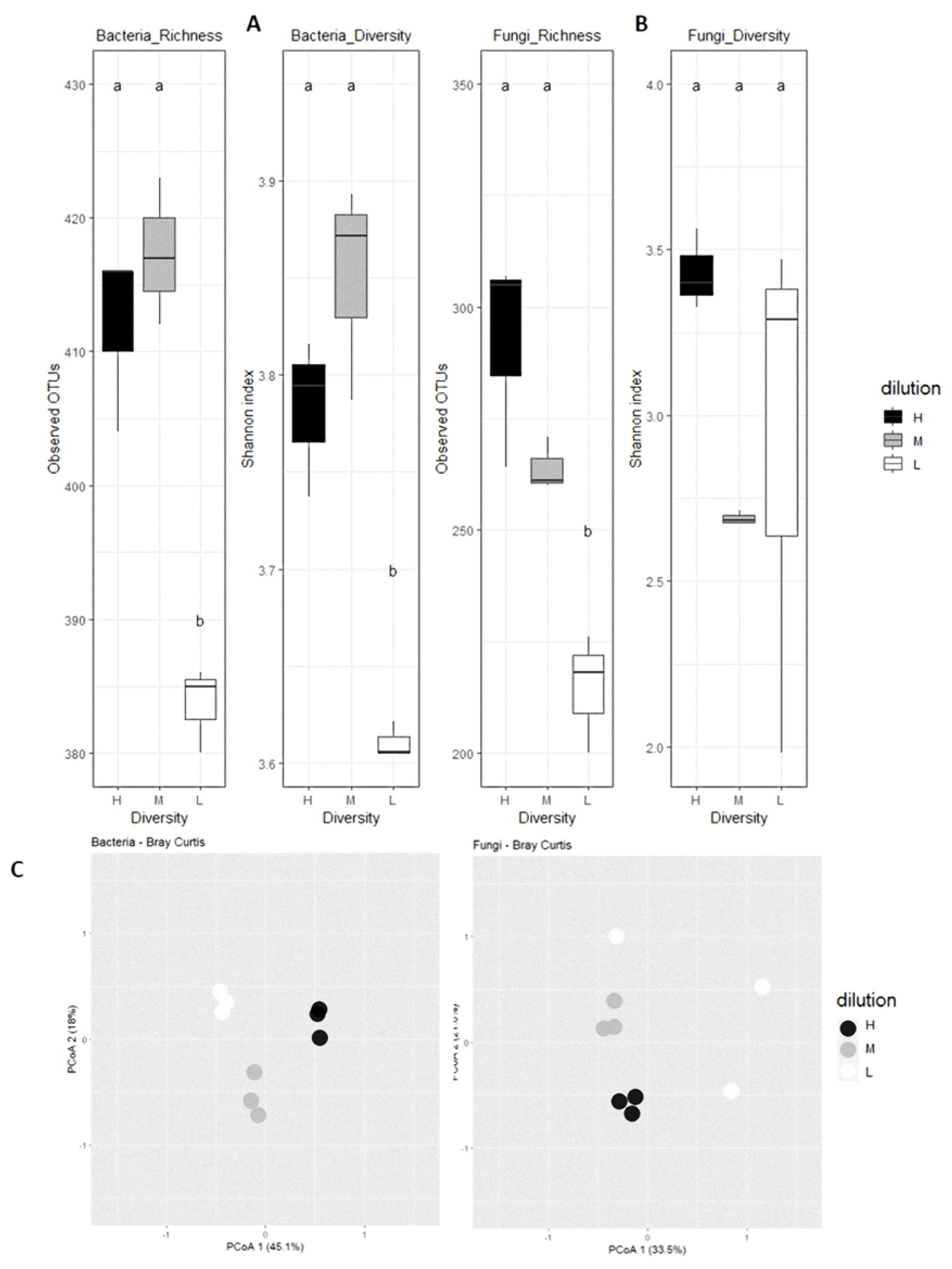
Bacterial (A) and fungal (B) richness and diversity, and communities’ structures (C) in the three soils used in this study. Mean richness (number of observed OTUs) and alpha-diversity (Shannon index) for the 3 soil microbial modalities (H, High in black; M, Medium in medium grey; L, Low in white) are presented in bacterial (A) and fungal (B) communities. Different letters indicate statistically significant differences among communities at P < 0.05. Principal coordinates analysis (PCoA) projection of the communities’ structure is shown for bacteria and fungi for the H, M and L diversities (C).

In conclusion, the soils obtained by microbial diversity manipulation through serial dilutions displayed different decreasing microbe richness and diversity, validating thus their use for evaluating their effect on *B. napus* infection by *P. brassicae*.

### Modulation of the plant susceptibility to clubroot according to the soil microbiota composition

The dry aerial parts were weighted in all experimental conditions (Fig 2A). At Ti (intermediary time), no significant differences were measured between healthy and inoculated plants, whatever both the soil microbiota modality and the plant genotype (except a small difference in H between healthy and inoculated Yudal). On the contrary, at the final time of the experiment (Tf), the inoculated plants displayed significant reduced aerial dry weight than healthy plants, whatever both the soil microbiota modality and the host plant genotype. At this time-point, the weight of aerial parts of both healthy and inoculated Tenor plants was weaker than in Yudal plants.

**Fig 2.**
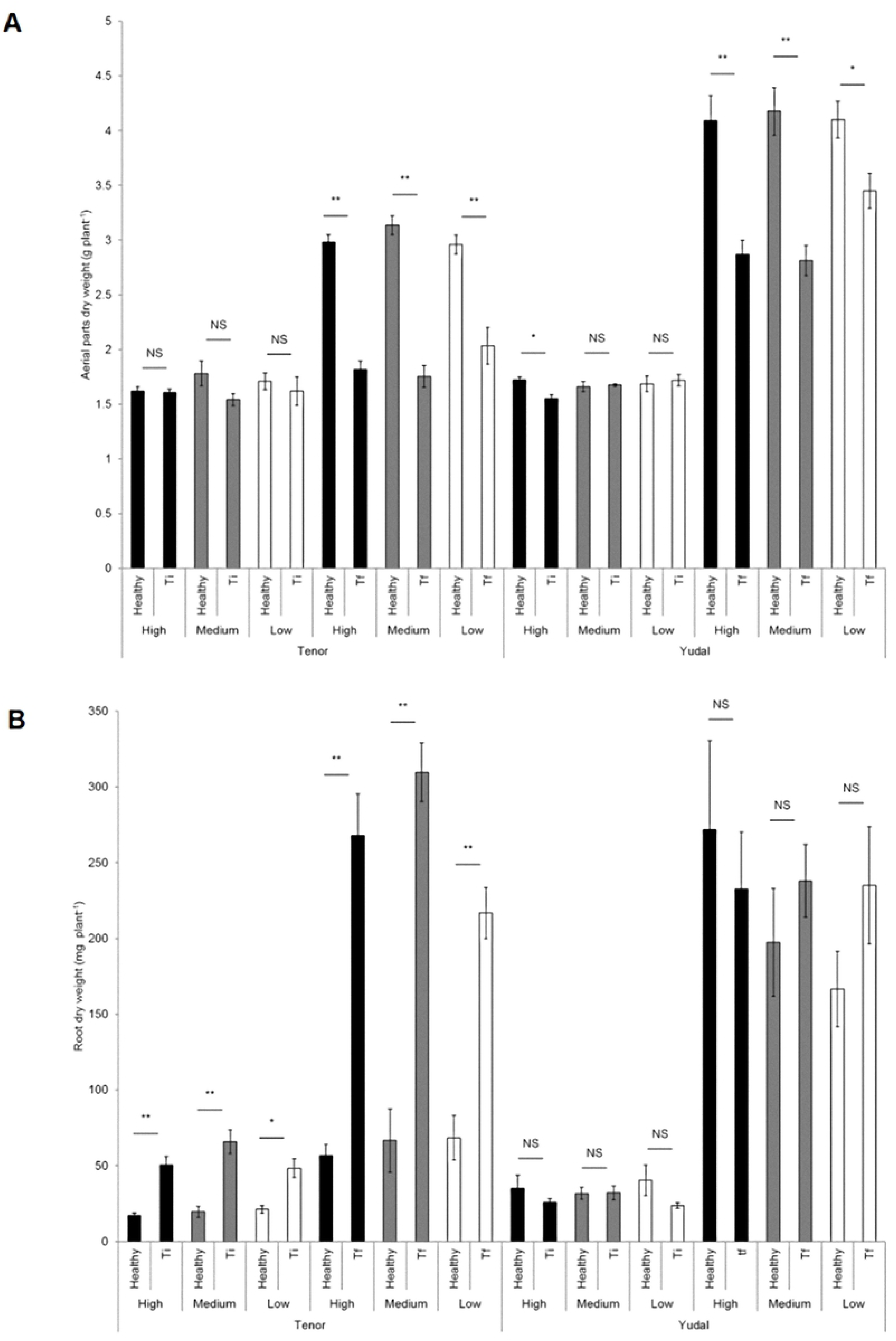
Aerial and root biomasses. The dry aerial parts (A) and roots (B) were weighted for both genotypes (Tenor and Yudal) at different days after inoculation (Ti, 28 dai; Tf 36 or 48 dai). For soil diversity, black, medium grey and white bars correspond to High (H), Medium (M) and Low (L) diversities, respectively. Error bars represent standard errors from the means of 8 plants. **, P < 0.01; *, P < 0.05; NS, Non Significant.

Concerning the roots (Fig 2B), the Tenor inoculated roots showed heavier dry mass (5 to 6 times more) at Ti and Tf than healthy roots, for each soil microbiota modality. The Tenor healthy roots had weak growth between Ti and Tf, whatever the soil, whereas inoculated Tenor had roots 6 times heavier at Tf than at Ti. This is the result of a strong development of galls in this genotype during this period. Concerning the Yudal root dry weights, no differences between healthy and infected plants were measured whatever the microbiota soil dilution and whatever the sampling date, probably because of the small size of galls clearly visible in Yudal genotype. At Tf, Yudal healthy roots were heavier than Tenor ones because of different root developmental patterns between the two genotypes.

At each sampling time, the soil microbiota modality had overall no effect on both aerial and root dry weights of healthy and inoculated plants.

At Ti and Tf, disease severity of inoculated plants was scored by determining the disease index (DI) and the DNA pathogen content (Fig 3). For each plant genotype, the DI showed the progression of disease along time-points: DI is about 50% at Ti and 80 % at Tf for Tenor, and less than 20% at Ti and 50% at Tf for Yudal. Whatever the soil modality and the sampling date, Yudal displayed lower DI than Tenor. This expected difference is consistent with the known level of clubroot resistance/susceptibility already described for these genotypes [64]. The soil microbiota modality had an effect on DI. For Tenor, at Ti and Tf, the DI was statistically significantly lower in L compared to H and M, and the highest DI was obtained in M. The DNA pathogen content followed the same pattern. At Ti, the *P. brassicae* DNA content was low, making difficult to compare the values between samples. At Tf, the DNA pathogen content was lower in L than in H and M, and higher in M, providing a bell-curve. Concerning the Yudal genotype, very low DI and DNA *P. brassicae* content were observed at Ti, making difficult the interpretation of the results. At Tf, decreasing gradients of DI and pathogen DNA content were measured through soil dilutions from H to L: the less rich and diverse soil, the less plant disease and DNA pathogen content.

**Fig 3.**
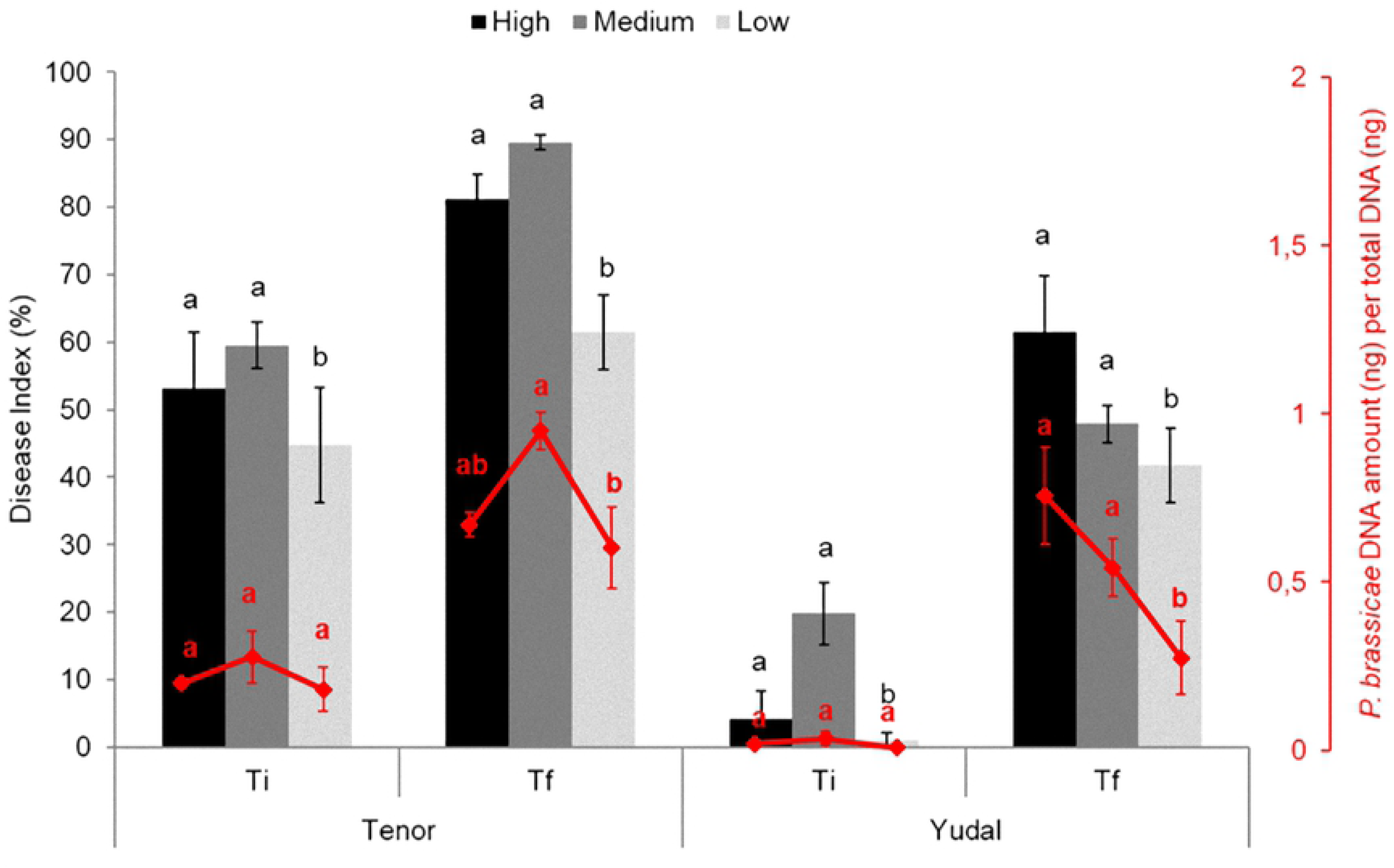
Influence of soil microbiota diversity on clubroot development. Plants were exposed to High (black), Medium (grey) or Low (white) soil microbial modalities during 28 (Ti), 36 or 48 (Tf) days after inoculation with the eH isolate of *P. brassicae*. The clubroot symptoms were estimated according to the disease index and the quantification of *P. brassicae* DNA by qPCR, expressed as a ratio of the 18S DNA quantity relative to the total DNA. Data are means of 3 biological replicates (12 plants per replicate) and error bars represent standard errors of the means. Means with different letters are statistically significantly different according to the analysis of variance test (P < 0.05).

### Overview, mapping and validation of RNAseq data

Approximately 80 to 100 million (M) reads by sample were obtained, and from 86 to 93% of the reads were mapped to the reference genome that we constructed, corresponding to the *B. napus* and the *P. brassicae* concatenated genomes.

Pathogen gene expression’s profiles were clearly clustered by the host plant genotype at Ti, and both by the soil microbiota modality and the host plant genotype at Tf (S3A Fig). No similar heatmap was performed with the *B. napus* gene expression profiles because of a huge number of expressed genes making the figure unreadable.

Hierarchical Cluster Analysis (HCA) (S3B Fig) on the filtered and normalized counts values concerning *P. brassicae* for each sample at Ti showed no true cluster structure in function of replicate, soil microbiota diversity or host plant genotype. On the contrary, at Tf for both host genotypes, the HCA analysis identified separated groups for the three replicates in H, in a lesser extent in M, and a less good grouping in L. This indicated that the experimental variation was higher in the more diluted soil microbial modality (L). Concerning the *B. napus* reads, in healthy (S4A Fig) and inoculated (S4B Fig) plants, the analysis showed that data clustered first by the host genotype, and then by the time factor, the soil modality and the replicate.

### Modulation of the *P. brassicae* transcriptome by the soil microbiota composition

Table 1 shows the number of DEGs in *P. brassicae* and *B. napus* according to H compared to M or L, for each inoculated host genotype. The comparisons are focused on differences between modalities considered closest to the initial state of the soil (*i.e.* H) and the diluted conditions (M and L).

**Table 1.**
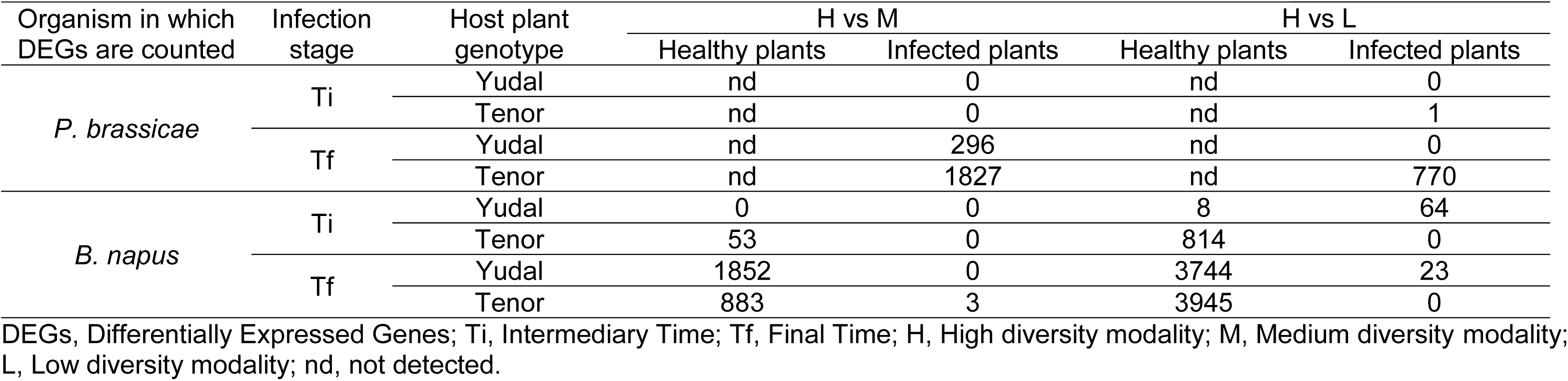
Number of DEGs in *P. brassicae* and in *B. napus* depending on the soil microbiota diversity levels.

Concerning the *P. brassicae* transcriptome, no DEGs between the soil microbiota modalities were detected at Ti (except only one gene between H and L in infected Tenor). On the contrary, at Tf, when galls were developed, the transcriptome of *P. brassicae* was different between soils. Interestingly, *P. brassicae* displayed a higher number of DEGs when infecting Tenor (2597 DEGs between H and both M and L) than when infecting Yudal (296 DEGs).

#### Modulation of the *P. brassicae* transcriptome by the soil microbiota composition when infecting Yudal

In the interaction with Yudal, only the M condition had an effect on the *P. brassicae* gene expression compared to H at Tf (Table 1). The complete list of the DEGs is presented in the S2 Table. Only nine genes among the 296 DEGs were overexpressed at M compared to H, with a small fold-change between conditions (1.2 to 1.6). No particular function of these genes can be easily associated with the DI between M and H (general pathways, such as signalization and chromosome condensation). On the contrary, a higher number of *P. brassicae* genes (287) were significantly underexpressed at M compared to H, in the same way than level of disease was lower at M compared to H. We selected the top 30 most significant down-regulated genes in M compared to H, with a fold-change greater than 2 (Table 2). Some of these top genes are potentially involved into the transport of molecules (e.g. *FMN-binding glutamate synthase family*, *MFS transporter Major Facilitator Superfamily*), and in development, growth and cell differentiation (e.g. *Chitin Synthase_2*, *Phosphoenolpyruvate carboxykinase*, *Glycosyltransferase*). Other genes were related to pathogenicity, including *Carbohydrate-binding module family_18*, *Glycoside hydrolase family_16*, and *NUDIX_hydrolase*.

**Table 2.** Selection of top 30 ranking *P. brassicae* highly down-regulated genes (fold-change > 2) in M compared to H at Tf when infecting Yudal (Y).

| *P. brassicae* gene | *P. brassicae* gene expression level |  | Fold-change | Description | Enzyme Codes |
| --- | --- | --- | --- | --- | --- |
|  | in Y / H / Tf | in Y / M / Tf |  |  |  |
| Pldbra_eH_r1s003g01588 | 0.43 | 0.02 | 11.42 | sugar_ABC_transporter_substrate-binding <sup>1</sup> | NA |
| Pldbra_eH_r1s004g02573 | 0.69 | 0.06 | 10.33 | ADP-ribosylation_factor_6 <sup>2</sup> | NA |
| Pldbra_eH_r1s003g01442 | 0.58 | 0.04 | 8.56 | calcium_calmodulin-dependent_kinase_type_IV-like <sup>2</sup> | ec:2.7.11.10 |
| Pldbra_eH_r1s003g01889 | 1.85 | 0.34 | 4.76 | NUDIX_hydrolase <sup>3</sup> | ec:3.6.1.65 |
| Pldbra_eH_r1s008g04734 | 2.32 | 0.46 | 4.72 | Serine_threonine-kinase_Sgk3 <sup>2</sup> | ec:3.1.4.4 |
| Pldbra_eH_r1s011g06165 | 4.32 | 0.89 | 4.53 | UDP-D-xylose:L-fucose_alpha-1;3-D-xylosyltransferase_1-like <sup>2</sup> | ec:2.4.1.37 |
| Pldbra_eH_r1s015g07579 | 6.72 | 1.51 | 4.21 | MFS_transporter <sup>1</sup> | NA |
| Pldbra_eH_r1s001g00152 | 1.67 | 0.40 | 4.06 | carbohydrate-binding_module_family_18 <sup>3</sup> | ec:3.2.1.14 |
| Pldbra_eH_r1s001g00029 | 7.63 | 1.84 | 4.02 | WD40_repeat | NA |
| Pldbra_eH_r1s008g04750 | 5.45 | 1.33 | 3.91 | methyltransferase_domain-containing | ec:2.1.1.300 |
| Pldbra_eH_r1s002g01072 | 7.08 | 1.78 | 3.89 | FMN-binding_glutamate_synthase_family <sup>1</sup> | ec:1.4.1.14 |
| Pldbra_eH_r1s001g00617 | 8.55 | 2.17 | 3.86 | glutamate_NAD(P)+ <sup>2</sup> | ec:1.4.1.23 |
| Pldbra_eH_r1s042g12180 | 3.67 | 0.94 | 3.75 | calcium/calmodulin-dependent_protein_kinase_type_IV-like <sup>2</sup> | ec:2.7.11.10 |
| Pldbra_eH_r1s007g04295 | 21.71 | 6.20 | 3.47 | Mps1_binder <sup>2</sup> | NA |
| Pldbra_eH_r1s001g00511 | 40.71 | 11.71 | 3.46 | serine_threonine-kinase_HT1 <sup>2</sup> | ec:2.7.11.10 |
| Pldbra_eH_r1s016g07781 | 26.37 | 7.58 | 3.45 | chitin_synthase_2 <sup>2</sup> | ec:2.4.1.16 |
| Pldbra_eH_r1s034g11599 | 8.64 | 2.45 | 3.41 | WD-40_repeat_domain-containing | NA |
| Pldbra_eH_r1s025g10321 | 8.55 | 2.46 | 3.38 | maltose_maltodextrin_ABC_substrate_binding_periplasmic <sup>1</sup> | ec:2.5.1.2 |
| Pldbra_eH_r1s014g07095 | 3.27 | 0.92 | 3.35 | glucosamine_6-phosphate_N-acetyltransferase <sup>2</sup> | ec:2.3.1.193 |
| Pldbra_eH_r1s001g00671 | 17.87 | 5.47 | 3.23 | glutathione-disulfide_reductase | ec:1.8.1.7;ec:1.8.2.3;ec:1.8.1.5 |
| Pldbra_eH_r1s003g01550 | 8.87 | 2.75 | 3.15 | glycosyltransferase <sup>2</sup> | ec:2.4.2.38 |
| Pldbra_eH_r1s003g01890 | 49.53 | 15.74 | 3.13 | glycoside_hydrolase_family_16 <sup>3</sup> | ec:3.2.1.151 |
| Pldbra_eH_r1s033g11505 | 14.72 | 4.86 | 3.00 | glycosyltransferase <sup>2</sup> | ec:2.4.2.38 |
| Pldbra_eH_r1s028g10813 | 7.61 | 2.63 | 2.80 | ABC_transporter_G_family <sup>1</sup> | ec:3.6.1.15;ec:3.6.3.43 |
| Pldbra_eH_r1s022g09622 | 47.16 | 17.11 | 2.74 | phosphoenolpyruvate_carboxykinase <sup>2</sup> | ec:4.1.1 |
| Pldbra_eH_r1s024g09958 | 32.88 | 12.19 | 2.68 | phosphate_ABC_transporter_substrate-binding <sup>1</sup> | ec:3.1.3.1 |
| Pldbra_eH_r1s002g00819 | 6.83 | 2.63 | 2.52 | cytochrome_P450 | ec:1.6.2.4;ec:1.14.14.1;ec:1.14.21.7;ec:1.16.1.5;ec:1.18.1.7 |
| Pldbra_eH_r1s028g10814 | 6.19 | 2.49 | 2.41 | ABC_transporter <sup>1</sup> | ec:3.6.1.3;ec:3.6.1.15;ec:3.6.3.43 |
| Pldbra_eH_r1s009g05056 | 9.63 | 4.23 | 2.28 | probable_phospholipid-transporting_ATPase_IA_isoform_X1 <sup>1</sup> | ec:3.6.1 |
| Pldbra_eH_r1s003g01928 | 56.23 | 27.59 | 2.03 | chitin_synthase_2 <sup>2</sup> | ec:2.4.1.16 |

Genes potentially involved in transport of molecules <sup>1</sup>, development and growth <sup>2</sup>, or pathogenicity <sup>3</sup>.

#### Modulation of the *P. brassicae* transcriptome by the soil microbiota composition when infecting Tenor

In the interaction with Tenor, 1827 genes of *P. brassicae* were differentially expressed at Tf between M and H (Table 1), most of them (1360 genes *i.e.* 75%) being overexpressed in M, and a smaller part (467 genes) underexpressed in M (S3A Table). Between L and H, there were 770 DEGs (S3B Table), with 532 *(i.e.* 70%) genes overexpressed in L compared to H and 238 underexpressed. In total, compared to the normal H level diversity, 621 *P. brassicae* genes were modulated both by M (out of 1827 genes, ie 34%) and L (out of 770 genes, ie 81%) conditions (S3C Table). Most of the genes regulated in L were also regulated in M. Moreover, these 621 genes displayed similar expression profiles: 450 genes were overexpressed at both M and L compared to H, and conversely for 171 genes. For these 171 genes, the fold-change was very small (< 1.5 for 169 genes whatever the comparison between soil microbiota diversities), but the gene expression levels were elevated. On the contrary, among the 450 genes overexpressed in M or L compared to H, 346 displayed a fold-change sharply higher than 2. The Table 3 shows the top 50 ranking by fold-change genes among these 346 *P. brassicae* genes overexpressed in M and L compared to H. Many of them were related to functions of transport (*phospholipid-transporting ATPase*, *FMN-binding_glutamate synthase*, *Ammonium transporter*, *Phosphate ABC_transporter* or *Potassium transporter*), growth (*Chitin synthase_2*), detoxification (*Glutathione_S transferase*, *Zinc_C2H2_type_family*), or potential pathogenicity (*E3-Ubiquitin ligase*, *alkaline ceramidase*, *cytosolic carboxypeptidase_4*, *serine carboxypeptidase_CPVL*).

**Table 3.**
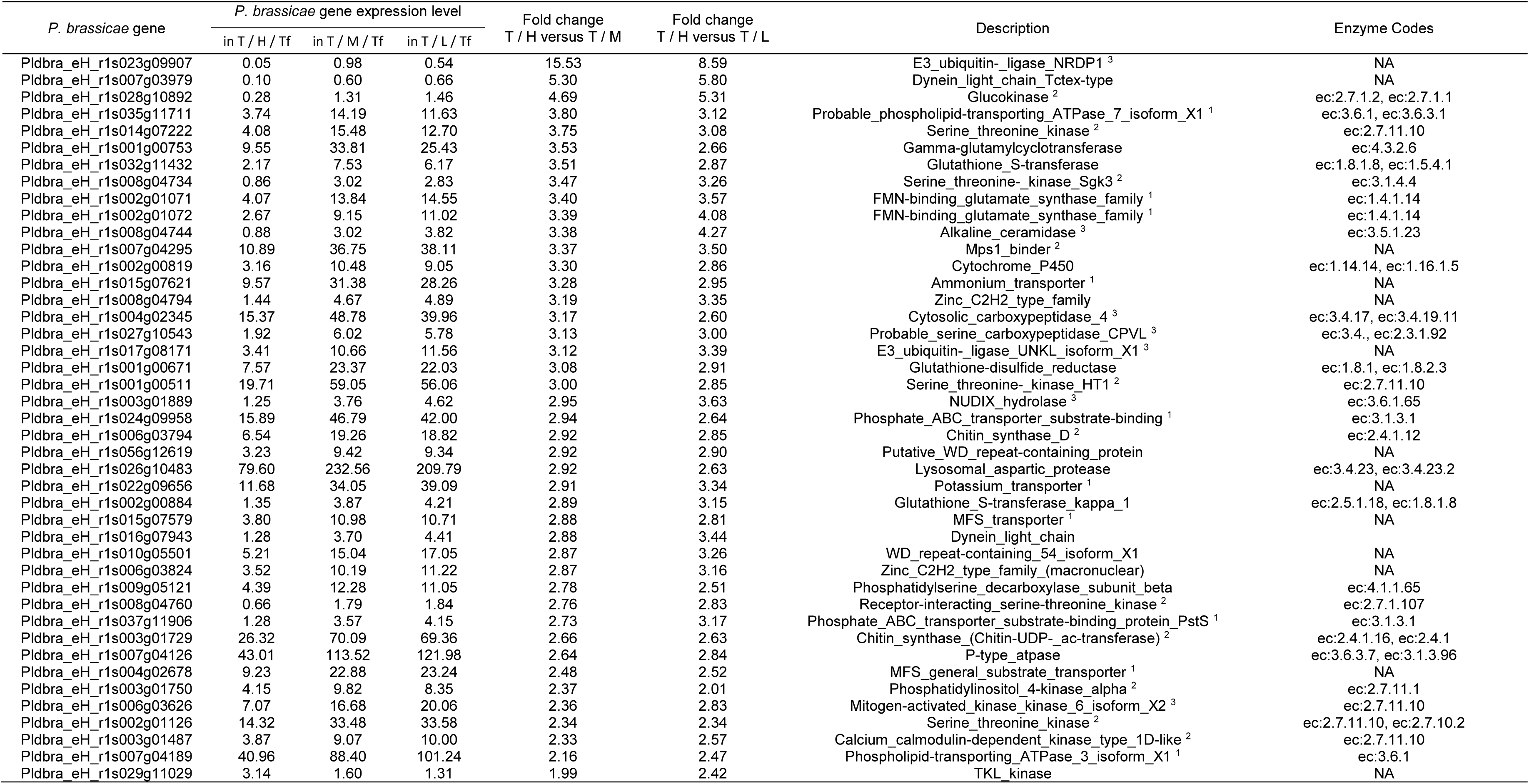
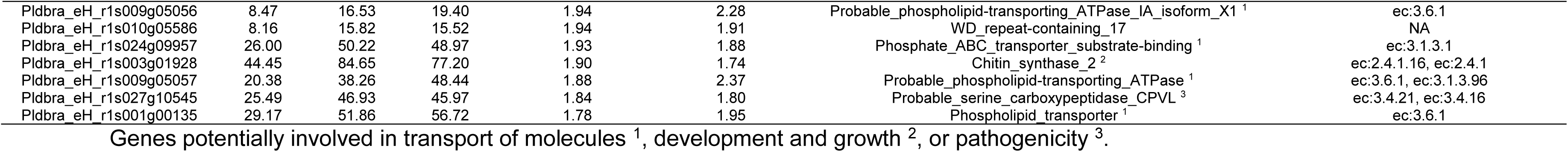
Selection of top 50 *P. brassicae* genes significantly differentially overexpressed in both M and L compared to H at Tf when infecting Tenor (T).

#### Focus on modulation of the *P. brassicae* transcriptome by the soil microbiota composition between H and M

We focused on the analyses of the *P. brassicae* gene expression between M and H at Tf because in these two soil microbiota modalities, we observed (i) the most important differences in pathogen gene expression for both plant genotypes, and (ii) a contrasted disease phenotype in function of the host plant genotype (Fig 3): lower disease level in M versus H in Yudal and higher disease level in M versus H in Tenor.

The sense of over- or under-expression profiles depending on the soil condition (H or M) was studied in detail in function of the host genotype. As shown in the Venn diagram (Fig 4), 1360 *P. brassicae* genes (out of 1827, *i.e.* 74%) when infecting Tenor, and only 9 *P. brassicae* genes (out of 296, *i.e.* 3%) when infecting Yudal were overexpressed in M compared to H. On the contrary, almost all the genes that were regulated by the soil microbiota diversity when Yudal was infected (260 out of 296) were underexpressed in M compared to H, although they were overexpressed in M versus H when infecting Tenor. The complete list of these 260 genes with the particular expression profile depending on the H / M levels and the host plant genotypes is indicated in the S4 Table. Among these 260 genes, a selection of the top 40 genes ranked according to the fold-change (Table 4) showed that the main functions encoded by these genes were related to the transport of molecules, the growth and development, the detoxification process and the pathogenicity. Concerning the 1100 genes specifically overexpressed in the Tenor genotype in L compared to H, most of them were related to transport of molecules (data not shown).

**Fig 4.**
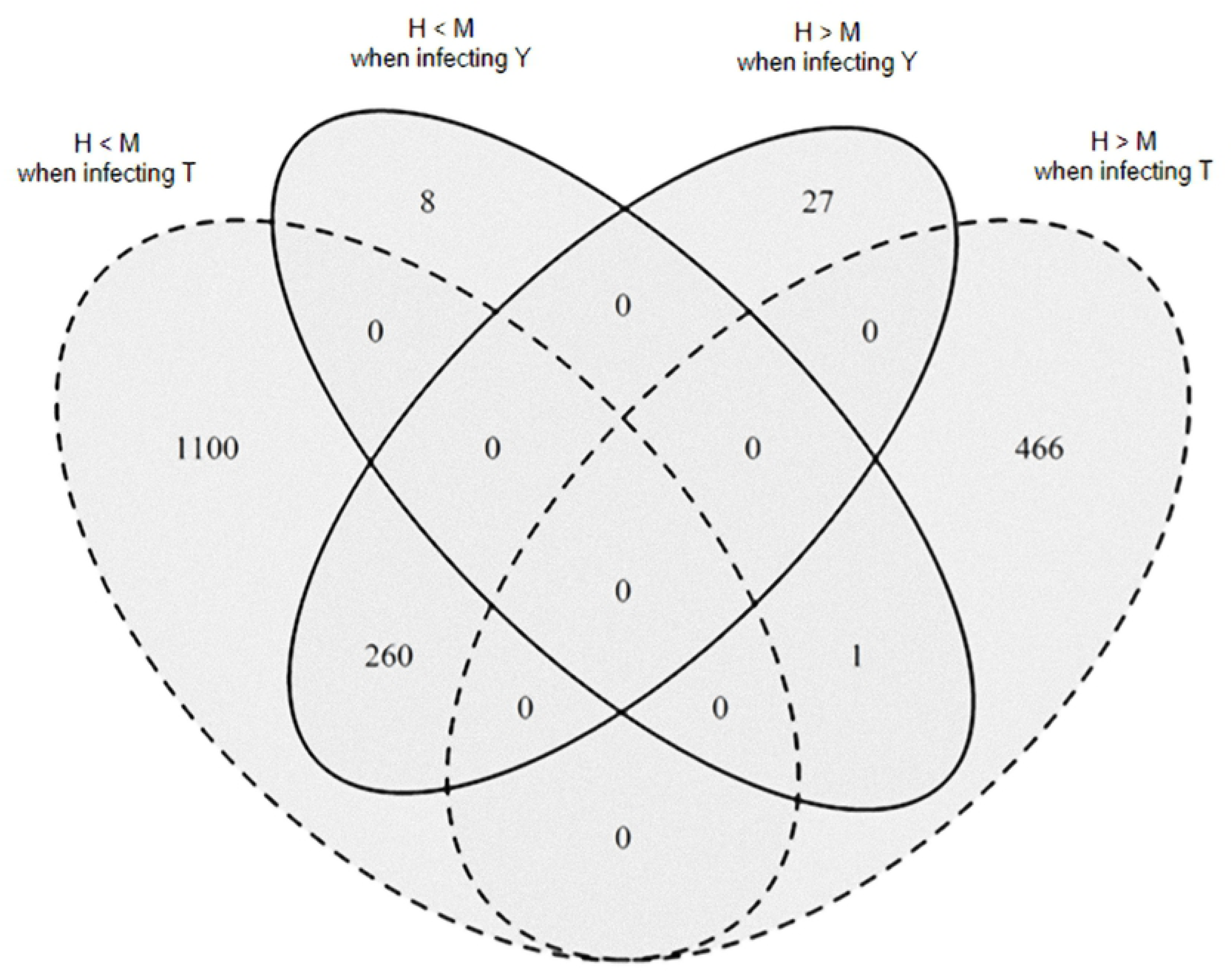
Number of *P. brassicae* differentially expressed genes (DEGs) at Tf between High (H) and Medium (M) soil microbial diversity levels when infected Yudal or Tenor. The Venn diagram shows the number of significantly *P. brassicae* DEGs (P < 0.05) that are overexpressed (M > H) or underexpressed (M < H) in M compared to H according to the host *B. napus* genotypes (Yudal, Y; Tenor, T) at the sampling date Tf.

**Table 4.** Selection of top 40 *P. brassicae* differentially expressed genes between H and M at Tf in an opposite sense when infecting Yudal (Y) or Tenor (T).

| *P. brassicae* gene | *P. brassicae* gene expression level |  | Fold change<br>Y / H versus Y / M | *P. brassicae* gene expression level |  | Fold change<br>T / H versus T / M | Description |
| --- | --- | --- | --- | --- | --- | --- | --- |
|  | in Y / H / Tf | in Y / M / Tf |  | in T / H / Tf | in T / M / Tf |  |  |
| Pldbra_eH_r1s001g00029 | 7.63 | 1.84 | 4.02 | 4.46 | 12.03 | 2.70 | WD40_repeat |
| Pldbra_eH_r1s001g00152 | 1.67 | 0.40 | 4.06 | 1.16 | 2.79 | 2.38 | carbohydrate-binding_module_family_18<sup>3</sup> |
| Pldbra_eH_r1s001g00179 | 6.45 | 2.15 | 2.90 | 2.86 | 9.67 | 3.35 | adenylate_guanylate_cyclase<sup>3</sup> |
| Pldbra_eH_r1s001g00511 | 40.71 | 11.71 | 3.46 | 19.71 | 59.05 | 3.00 | Serine_threonine-kinase_HT1<sup>2</sup> |
| Pldbra_eH_r1s001g00617 | 8.55 | 2.17 | 3.86 | 4.59 | 11.96 | 2.60 | glutamate_NAD(P)+<sup>2</sup> |
| Pldbra_eH_r1s001g00671 | 17.87 | 5.47 | 3.23 | 7.57 | 23.37 | 3.08 | glutathione-disulfide_reductase<sup>4</sup> |
| Pldbra_eH_r1s001g00753 | 42.72 | 16.10 | 2.64 | 9.55 | 33.81 | 3.53 | gamma-glutamylcyclotransferase<sup>4</sup> |
| Pldbra_eH_r1s002g00819 | 6.83 | 2.63 | 2.52 | 3.16 | 10.48 | 3.30 | cytochrome_P450<sup>4</sup> |
| Pldbra_eH_r1s002g00884 | 1.82 | 0.48 | 3.42 | 1.35 | 3.87 | 2.89 | glutathione_S-transferase_kappa_1_[Rhodotorula_toruloides_NP11]<sup>4</sup> |
| Pldbra_eH_r1s002g01072 | 7.08 | 1.78 | 3.89 | 2.67 | 9.15 | 3.39 | FMN-binding_glutamate_synthase_family<sup>1</sup> |
| Pldbra_eH_r1s003g01442 | 0.58 | 0.04 | 8.56 | 0.29 | 1.19 | 4.01 | calcium_calmodulin-dependent_kinase_type_IV-like<sup>2</sup> |
| Pldbra_eH_r1s003g01550 | 8.87 | 2.75 | 3.15 | 6.10 | 16.04 | 2.62 | Glycosyltransferase_uncharacterized<sup>2</sup> |
| Pldbra_eH_r1s003g01889 | 1.85 | 0.34 | 4.76 | 1.25 | 3.76 | 2.95 | NUDIX_hydrolase<sup>3</sup> |
| Pldbra_eH_r1s003g01890 | 49.53 | 15.74 | 3.13 | 26.18 | 68.19 | 2.60 | glycoside_hydrolase_family_16<sup>3</sup> |
| Pldbra_eH_r1s003g01928 | 56.23 | 27.59 | 2.03 | 44.45 | 84.65 | 1.90 | chitin_synthase_2<sup>2</sup> |
| Pldbra_eH_r1s006g03824 | 6.92 | 2.18 | 3.10 | 3.52 | 10.19 | 2.87 | Zinc_C2H2_type_family_(macronuclear)<sup>4</sup> |
| Pldbra_eH_r1s007g04126 | 75.85 | 26.51 | 2.85 | 43.01 | 113.52 | 2.64 | p-type_atpase |
| Pldbra_eH_r1s007g04295 | 21.71 | 6.20 | 3.47 | 10.89 | 36.75 | 3.37 | Mps1_binder<sup>2</sup> |
| Pldbra_eH_r1s008g04734 | 2.32 | 0.46 | 4.72 | 0.86 | 3.02 | 3.47 | Serine_threonine-kinase_Sgk3<sup>2</sup> |
| Pldbra_eH_r1s008g04750 | 5.45 | 1.33 | 3.91 | 3.65 | 10.15 | 2.78 | methyltransferase_domain-containing |
| Pldbra_eH_r1s008g04794 | 3.37 | 0.84 | 3.69 | 1.44 | 4.67 | 3.19 | zinc_C2H2_type_family<sup>4</sup> |
| Pldbra_eH_r1s009g05056 | 9.63 | 4.23 | 2.28 | 8.47 | 16.53 | 1.94 | probable_phospholipid-transporting_ATPase_IA_isoform_X1<sup>1</sup> |
| Pldbra_eH_r1s009g05121 | 8.82 | 2.71 | 3.20 | 4.39 | 12.28 | 2.78 | phosphatidylserine_decarboxylase_subunit_beta |
| Pldbra_eH_r1s011g06165 | 4.32 | 0.89 | 4.53 | 2.28 | 7.67 | 3.34 | UDP-D-xylose:L-fucose_alpha-1, 3-D-xylosyltransferase_1-like<sup>2</sup> |
| Pldbra_eH_r1s014g07095 | 3.27 | 0.92 | 3.35 | 1.72 | 5.57 | 3.17 | glucosamine_6-phosphate_N-acetyltransferase<sup>2</sup> |
| Pldbra_eH_r1s015g07579 | 6.72 | 1.51 | 4.21 | 3.80 | 10.98 | 2.88 | MFS_transporter<sup>1</sup> |
| Pldbra_eH_r1s016g07781 | 26.37 | 7.58 | 3.45 | 13.19 | 38.91 | 2.95 | chitin_synthase_2<sup>2</sup> |
| Pldbra_eH_r1s022g09622 | 47.16 | 17.11 | 2.74 | 22.80 | 62.09 | 2.72 | phosphoenolpyruvate_carboxykinase<sup>2</sup> |
| Pldbra_eH_r1s022g09656 | 23.28 | 7.19 | 3.21 | 11.68 | 34.05 | 2.91 | potassium_transporter<sup>1</sup> |
| Pldbra_eH_r1s024g09958 | 32.88 | 12.19 | 2.68 | 15.89 | 46.79 | 2.94 | phosphate_ABC_transporter_substrate-binding<sup>1</sup> |
| Pldbra_eH_r1s025g10321 | 8.55 | 2.46 | 3.38 | 4.50 | 10.67 | 2.38 | Maltose_maltodextrin_ABC_substrate_binding_periplasmic<sup>1</sup> |
| Pldbra_eH_r1s027g10543 | 2.67 | 0.79 | 2.96 | 1.92 | 6.02 | 3.13 | probable_serine_carboxypeptidase_CPVL<sup>3</sup> |
| Pldbra_eH_r1s028g10813 | 7.61 | 2.63 | 2.80 | 2.22 | 7.21 | 3.23 | ABC_transporter_G_family<sup>1</sup> |
| Pldbra_eH_r1s028g10814 | 6.19 | 2.49 | 2.41 | 1.74 | 6.07 | 3.42 | ABC_transporter<sup>1</sup> |
| Pldbra_eH_r1s033g11505 | 14.72 | 4.86 | 3.00 | 10.02 | 24.24 | 2.42 | Glycosyltransferase_uncharacterized<sup>2</sup> |
| Pldbra_eH_r1s034g11599 | 8.64 | 2.45 | 3.41 | 3.73 | 11.80 | 3.15 | WD-40_repeat_domain-containing |
| Pldbra_eH_r1s035g11711 | 7.18 | 2.74 | 2.57 | 3.74 | 14.19 | 3.80 | probable_phospholipid-transporting_ATPase_7_isoform_X1<sup>1</sup> |
| Pldbra_eH_r1s042g12180 | 3.67 | 0.94 | 3.75 | 1.55 | 5.79 | 3.66 | calcium/calmodulin-dependent_protein_kinase_type_IV-like<sup>2</sup> |
| Pldbra_eH_r1s056g12619 | 5.28 | 1.46 | 3.59 | 3.23 | 9.42 | 2.92 | putative_WD_repeat-containing_protein |
| Pldbra_eH_r1s058g12634 | 7.62 | 2.00 | 3.67 | 4.64 | 12.69 | 2.74 | peptidase_M14 |

Genes potentially involved in transport of molecules<sup>1</sup>, development and growth<sup>2</sup>, pathogenicity<sup>3</sup>, or detoxification<sup>4</sup>.

#### Modulation of the *P. brassicae* transcriptome by the host plant genotype in each condition of soil microbiota composition

The number of DEGs in *P. brassicae* according to the plant host genotype for each microbial diversity is presented in the Fig 5. At Ti, the effect of the host plant genotype on *P. brassicae* transcriptome was more important in H (445 DEGs) than M (2 DEGs) or L (60 DEGs), and most of the DEGs in L (78%) were also DEGs in H. Only one gene (with no known annotation) was differentially expressed according to the host genotype whatever the soil microbiota diversity. At Tf, a higher number of DEGs was found between host genotypes for each diversity than at Ti. The effect of the plant genotype was around 6 times more important in M (3896 DEGs) than in H (604 DEGs) or L (560 DEGs). This is coherent with the observation that the M condition led to a contrasted disease phenotype in function of the host plant genotype (Figure 3: higher disease level in H versus M for the infected Yudal and lower disease level in H versus M for the infected Tenor). There were only 31 common DEGs between H and L and 154 between H and M, showing a particular *P. brassicae* transcriptome in function of the plant genotype in H. On the contrary, most of the DEGs in L were also DEGs in M. Finally, 84% (3262 out of 3896) of the *P. brassicae* DEGs between host genotypes in M were specific of this soil microbiota diversity. A core of 28 DEGs was common to the three soil modalities; among them, whatever the soil microbiota diversity, 11 and 17 were under- or over-expressed in Tenor compared to Yudal, respectively. These genes displayed either unknown functions or functions of the general metabolism (data not shown).

**Fig 5.**
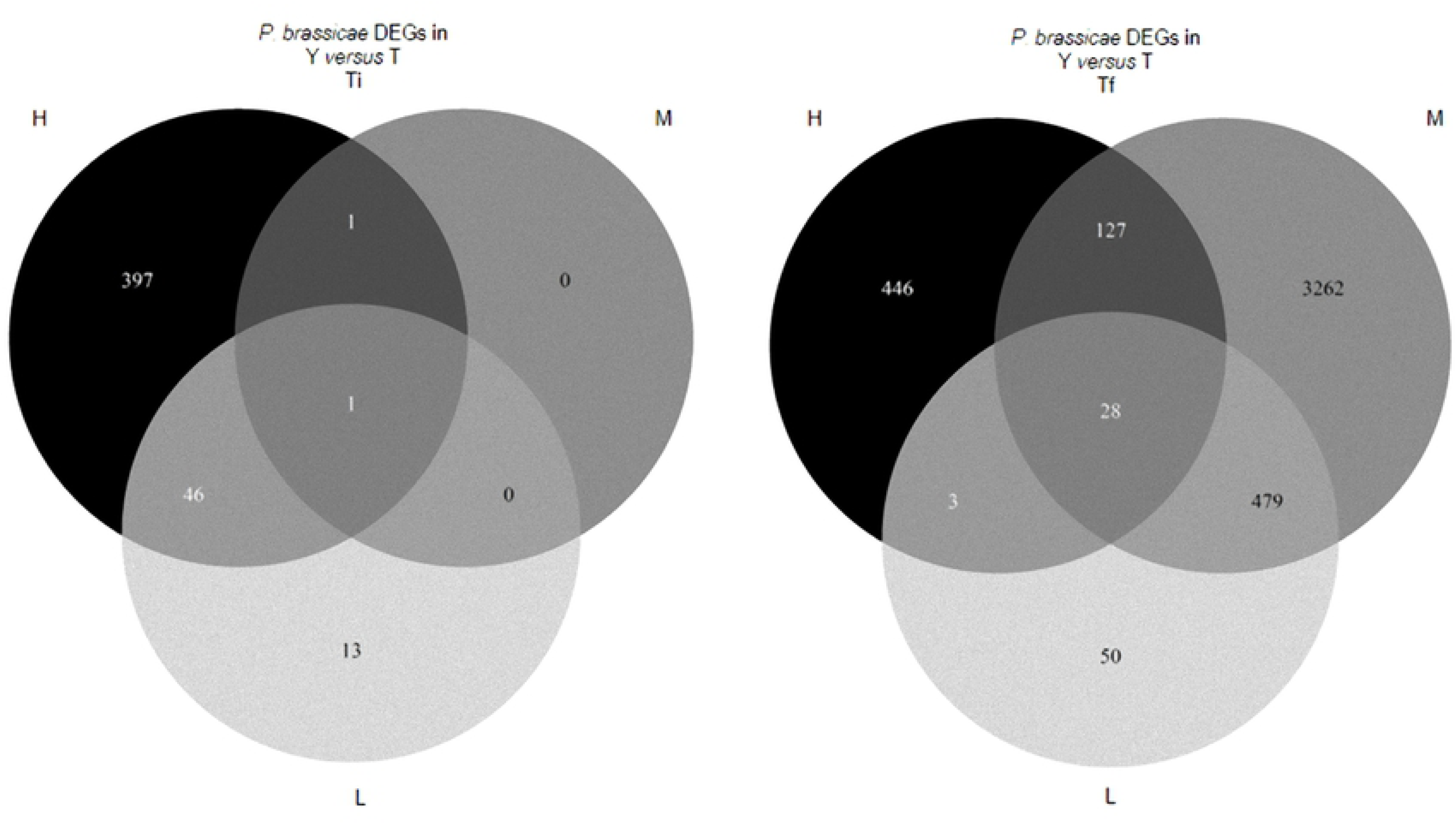
Number of *P. brassicae* differentially expressed genes (DEGs) in function of the host plant genotype for each soil microbial diversity level. The Venn diagram shows the number of significantly DEGs (P < 0.05) according to the host *B. napus* genotypes (T, Tenor; Y, Yudal) for each soil microbial diversity level (H, High; M, Medium; L, Low) at the sampling dates Ti and Tf.

### Modulation of the *B. napus* transcriptome by the soil microbiota composition

The results of soil diversity manipulation (M versus H and L versus H) at Ti and Tf on the *B. napus* transcriptome for each genotype, both in healthy and infected plants, are shown in the Table 1.

#### Modulation of the Yudal transcriptome by the soil microbiota composition

In healthy Yudal, a very moderate soil condition’s effect on DEGs number at Ti (0 to 8 genes), and a higher effect at Tf (1852 to 3744 genes) were measured.

In infected Yudal, the M condition did not modify the gene expression compared to H, although 64 genes at Ti (S5A Table) and 23 genes at Tf (S5B Table) were differentially expressed between L and H. Interestingly, the Yudal transcriptome was modified by L at Ti, although no effect of the diversity on plant disease phenotype was significantly detectable at this stage (Fig 3). At Tf, the number of the genes that were down/up-regulated was less than at Ti despite a more pronounced difference in disease phenotype between L and H. In Table 5 is shown a selection of *B. napus* genes for which the expression was greatly different in Yudal between L and H. The DEGs included a large number of genes encoding various proteins involved in plant defense, and particularly in hormonal pathways.

**Table 5.**
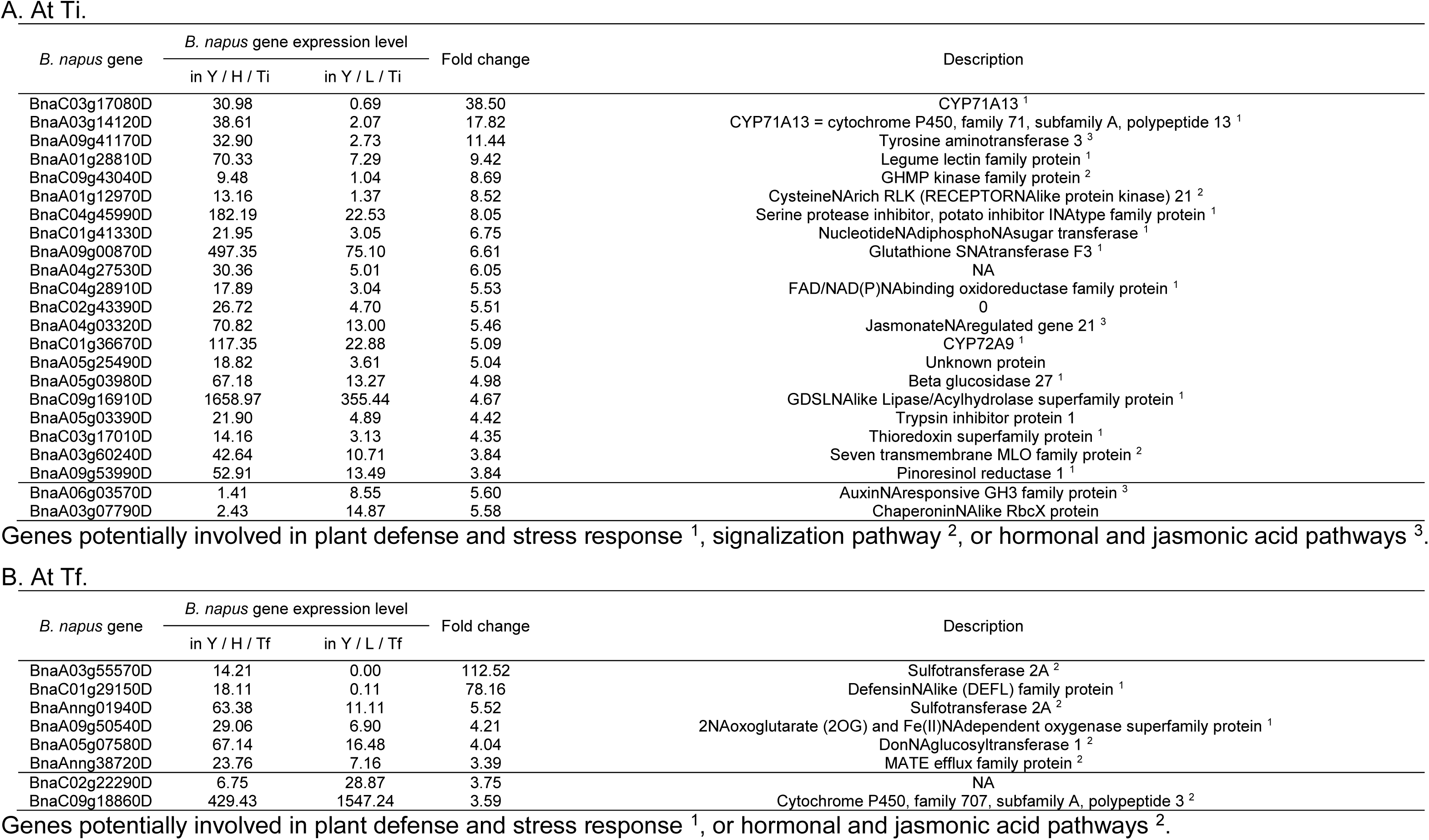
Selection of top Yudal differentially expressed genes between H and L at Ti (A) and Tf (B) when infected by *P. brassicae*.

#### Modulation of the Tenor transcriptome by the soil microbiota composition

In healthy Tenor, similar expression profiles to those of healthy Yudal were found, with a moderate number of DEGs at Ti between M and H (53 genes), and higher number between L and H (814 corresponding nearly to only 8 ‰ of the total number of expressed genes in *B. napus*). At Tf, 883 DEGs between M and H, and 3945 between L and H were found. In infected Tenor, no genes were differentially expressed between the soil conditions, except only 3 genes between M and H at Tf.

#### Host plant genotype’s effect on the *B. napus* transcriptome in each modality of soil microbiota composition

The global view of DEGs in healthy and infected plants of the two host genotypes, according to the soil microbiota modality and the interaction time is illustrated in Venn diagrams (S5 Fig). The number of *B. napus* DEGs between genotypes was huge in healthy and infected plants, and largely the same whatever the soil microbiota (14,789 to 27,537). In all the studied conditions, the effect of the genotype on plant transcriptome was very marked since about one third of the genes was differentially expressed between genotypes whatever the diversity, the time of interaction and the presence or not of the pathogen.

#### Modulation of the *B. napus* transcriptome by the infection stage in each modality of soil microbiota composition

The number of *B. napus* DEGs in each soil microbiota condition according to the infection stage showed high changes in transcript levels (S6 Fig). The high number of DEGs was retrieved for both host plant genotypes, infected or not, and for the three soil conditions. Whatever the diversity of the soil microbial community, the number of DEGs was quite similar for both genotypes in healthy plants. In infected plants, the number of DEGs in Yudal was slightly higher than in Tenor, particularly in H (19230 and 13771 DEGs in Yudal and Tenor, respectively) and L (15560 and 10547 DEGs in Yudal and Tenor, respectively). Depending on the soil condition, both genotypes displayed 25 to 50% of common DEGs set between Ti and Tf. A moderate number of DEGs was shared between plant genotypes and soil microbiota diversities (1388 and 2192 in healthy and infected plants, respectively).

By focusing more specifically on the *B. napus* genes that were differentially expressed between Ti and Tf for both infected genotypes and for the three soil’s conditions, 2192 genes were recovered (S6 Fig). Most of them were regulated in the same sense for Yudal and Tenor according to the time-point (S7A Fig). A slight part of genes had opposite sense of expression between plant genotypes: 6 genes were underexpressed in Yudal at Ti compared to Tf but over-expressed in Tenor at Ti compared to Tf, and 34 genes were over-expressed in Yudal at Ti compared to Tf but underexpressed in tenor at Ti compared to Tf (S7A Fig). The annotation of 33 genes out of the 40 was retrieved (S7B Fig). Concerning the genes overexpressed in Yudal and underexpressed in Tenor at Ti compared to Tf, they were mainly related to growth and plant development. Other genes were related to the response to disease, or involved in hormonal signalization. Two genes (*WRKY DNA binding protein 11* and *Basic region/leucine zipper motif 53*) encoding for transcription factors were also differentially expressed between Ti and Tf in a different way according to the plant genotype.

## Discussion

The plant-associated microbiota is more and more recognized as important determinant of plant health and pathogen suppression. As main ways to control clubroot such as crop rotations and cultivation of varieties carrying major resistance genes [29, 65] have shown their limits, there is a need to design alternative and durable methods based on ecological concepts. Exploring and understanding the mechanisms of disease regulation by microbiota could contribute to the emergence of innovative plant protection strategies.

Our research provides an extensive study of molecular mechanisms involved in complex host-pathogen interactions modulated by soil microbiota composition, using dual RNA-Seq to simultaneously capture the transcriptome of the two interacting partners. This approach has been applied to investigate a variety of host-pathogen relationships in major plant diseases in simplified in vitro experiments [66–68]. Our study upgraded the dual RNA-seq approach in more complex and realistic interaction’s conditions.

### Soil microbiota composition and clubroot phenotypes

The soil microbial diversity manipulation through serial dilutions (‘dilution to extinction’ experiment) led to a decreasing gradient of bacterial and fungal richness and a modification community’ structure, as previously described [25], allowing controlled experiments using different microbial diversity reservoirs with common soil properties. We found that the microbial diversity modulated the clubroot development, in different patterns according to the host plant genotype. Interestingly, when Yudal was infected, the decrease in microbial diversity led to a proportional decrease in disease level, and in infected Tenor, a bell curve of disease level according to microbial diversity was found. The invasion of pathogens is often described as linked to the level of microbial community’s diversity and connectedness [69, 70]. It is also known that rhizosphere and endophytic microbial communities, that play key roles in controlling pathogens [18, 27, 71, 72], are recruited from the communities of microorganisms in the soil in part in a plant-specific controlled way. It is indeed proved that different genotypes of the same plant species may have significant impacts on selecting rhizospheric partners through production of diverse root exudates [16, 73]. For instance, root-associated microbiota displaying reproducible plant genotype associations was recently identified in maize [74]. Genotype effects of the plant hosts can be also more important for individual microbial species [75]. The difference in modulation of clubroot by the soil microbial diversity between Yudal and Tenor, as well as the higher changes in *P. brassicae* transcript levels in function of soil microbiota composition when Tenor was infected compared to Yudal, could be due to a plant genotype’s effect on the process of microbial recruitment. More particularly, missing microbes, or prevalence of ‘helper’ microbes, or changes in the strength and connection of the microbes’ network between H, M or L conditions can support the disease’s outbreak [76]. Moreover, we previously showed that not only the structure of microbial communities associated with the rhizosphere and roots of healthy Brassica plants *(B. rapa)* evolved over time, but also that the invasion by *P. brassicae* changed root and rhizosphere microbial communities already assembled from the soil [28]. All these results highlighted the complexity of the microbial interactions in soil, including interactions between microorganisms, between microbes and plant, and between microbes and pathogen.

### Soil microbiota composition and *P. brassicae* transcriptome

The global view of distribution of DEGs according to the soil microbiota composition, in each plant genotype and time-point, showed that the *P. brassicae* transcriptome was not only more modulated when infected Tenor than Yudal, but also most strongly activated at Tf than Ti. During its life cycle, *P. brassicae* survives in soil in the form of resting spores. Sensing signal molecules, such as host root exudate production or specific soil environment, is essential to exit dormancy, trigger germination and begin the initial step of the life cycle inside the root: at this stage, suitable conditions in environment, such as the soil microbial diversity and composition, are necessary. Bi et al. [35] showed that *P. brassicae* is able to have perception of external signals thanks to specific signaling pathway and to adapt to its environment. In our study, the very early step of interaction between *P. brassicae* spores and soil microbiota was not measured. But the higher *P. brassicae* transcriptome modulation at Tf than at Ti highlighted the secondary cortical infection stage of clubroot disease as crucial for interaction between *P. brassicae* and the microbiota. In the same way, the root and rhizosphere-associated community assemblies in *B. rapa*, particularly the endophytic bacterial communities, were also strongly modified by *P*. *brassicae* infection during this stage [28]. Thus, the disturbance consequences of the interactions between *P*. *brassicae* and the endophytic communities inside the roots occurred at the tardive date of sampling, and the effect of soil environment on *P. brassicae* transcriptome was thereby measurable at the stages where the pathogen was in a close interaction with its host.

#### The soil microbiota composition affects the expression of *P. brassicae* genes potentially involved in the transport of molecules

At Tf, higher *P. brassicae* amount (and DI) were found in H compared to M in infected Yudal, whereas lower in H compared to M when infected Tenor. The DEGs in this same sense as *P. brassicae* amount between H and M were particularly analyzed for both infected host plant genotypes (Tables 2, 3, 4), and studied in function of their potential involvement in different functions. This is for example the case for several genes, overexpressed in conditions where DNA *P. brassicae* content was higher, that were related to functions of molecule transport. The loss of key biosynthetic pathways is indeed a common feature of parasitic protists, making them heavily dependent on scavenging nutrients from their hosts. Salvage of nutrients by parasitic protists is often mediated by specialized transporter proteins that ensure the nutritional requirements. This is the case of genes coding for a FMN-binding_glutamate_synthase, a complex iron–sulfur flavoprotein that plays a key role in the ammonia assimilation pathways also found in bacteria, fungi and plants [77, 78], and for a phospholipid transporting ATPase, a Phosphate_ABC_transporter or a Potassium transporter. Some transporters, such as the Ammonium_transporters are also expressed during host colonization and pathogenicity in fungus because of the importance of ammonia in host alkalinization [79, 80]. The soil microbiota composition and then the subsequent recruitment of endophyte microbes by the plant could affect the *P. brassicae* ability to recruit nutriments from the host because of potential competition for resource [81].

#### The soil microbiota composition affects the expression of *P. brassicae* genes potentially involved in growth and development

Other examples of DEGs between soil microbial diversities with expression profiles correlated to clubroot development were related to functions of growth, development and cell differentiation. For instance, the gene coding for a Chitin synthase, essential for the cell wall chitin depositions during resting spore maturation, was overexpressed in conditions where clubroot symptoms were more pronounced. The chitin-related enzymes are enriched in *P. brassicae* genome [32, 37, 39]. Deletion of *chitin synthase* genes in fungi most often results in developmental defects, which include defective infection structure development or defunct invasive growth [82, 83]. Concerning the gene coding for a Phosphoenolpyruvate_carboxykinase, its differential expression could make possible to *P. brassicae* a glucose-independent growth [84]. The differential expression of a gene coding for a Glycosyltransferase could facilitate the growth as shown in filamentous pathogenic fungi [85].

#### The soil microbiota composition affects the expression of *P. brassicae* genes potentially involved in pathogenicity

Some *P. brassicae* genes coding for potential pathogenicity factors, that were overexpressed in M compared to H in Tenor and/or underexpressed in M compared to H in Yudal, may explain in part the different disease phenotype observed in function of the soil microbial diversities’ conditions.

This was the case for the gene encoding a Glutathione transferase that was overexpressed in conditions of important clubroot development symptoms. Glutathione transferases represent an extended family of multifunctional proteins involved in detoxification processes and tolerance to oxidative stress. In *Alternaria brassicicola*, Glutathione transferases participate in cell tolerance to isothiocyanates, allowing the development of symptoms on host plant tissues [86]. The pathogenicity of *P. brassicae* could be partly related to its ability to protect itself against such plant defenses compounds.

For other genes putatively related to pathogenicity, we found the same trend of overexpression in conditions of important clubroot development. The E3-Ubiquitin ligase is described as a microbial effector protein that evolved the ability to interfere with the host E3-Ub-ligase proteins to promote disease [87]. The alkaline ceramidase is involved in the virulence of microbes like *Pseudomonas aeruginosa* [88]. The cytosolic carboxypeptidase_4 and the serine carboxypeptidase_CPVL are also described as potential factors of virulence with a role in adherence process, penetration of tissues, and interactions with the immune system of the infected host [89, 90]. The genes coding for the Carbohydrate-binding module_family_18 or the Glycoside_hydrolase family_16 can protect some fungi against plant defense mechanisms [91, 92]. For instance, CBM18-domain proteins protect from breakdown by chitinase in some fungi [83]. In Plasmodiophorids, proteins containing a CBM18 domain, could bind to the chitin in order to promote modification into chitosan, a weaker inducer of immune responses than chitin in many plants [32].

Finally, a conserved effector gene in the genomes of a broad range of phytopathogenic organisms across kingdoms (bacteria, oomycetes, fungi) [93, 94], the *NUDIX_hydrolase*, was found overexpressed in conditions where clubroot symptoms were highest, according to the soil microbial diversity. In *Arabidopsis thaliana* infected by *P. brassicae*, proteomics studies had already detected an upregulation of the NUDIX protein [95]. NUDIX effectors have been validated as pathogenesis players in a few host–pathogen systems, but their biological functions remain unclear [93]. Further studies are necessary to decipher if *P. brassicae* might share strategy involving NUDIX effectors described in other plant pathogens. The *NUDIX* gene is a good pathogenicity candidate gene, potentially responsible for *P. brassicae* infection and subsequent disease progression and that needs to be functionally assessed.

### Soil microbiota composition and *B. napus* transcriptome

#### The host plant genotype and the infection’s kinetic strongly affect the plant transcriptome whatever the soil microbiota composition

In both healthy and infected plants, the number of *B. napus* DEGs between genotypes was huge and largely shared between soil microbiota, and the number of plants DEGs between Ti and Tf was also high for each genotype whatever the soil microbiota composition. This demonstrates that the genetic control of the developmental process is highly dynamic and complex, and remains largely unknown.

The list of common DEGs between Ti and Tf in both genotypes and the three H, M, L conditions (S7 Fig) was studied more in detail, and particularly the genes overexpressed in Yudal but underexpressed in Tenor at Ti compared to Tf. These genes were mainly related to growth and plant development: *Sterol methyltransferase 3* [96], *C2H2like zinc finger protein* [97], *BES1/BZR1 homolog 2* [98], *WUSCHEL related homeobox 4* [99], *Expansin A1* [100], *Arabinogalactan protein 22* [101], *Trichome BireFringence 27* [102], *SKU5 similar 17* [103], *Transcription elongation factor (TFIIS) family protein* [104], *Endoxyloglucan transferase A3* [105], *KIPrelated protein 2* [106], and *Ras related small GTPNAbinding family protein* [107]. Other genes of the list were related the response to disease, like the *RING/box superfamily protein* (*family E3 ligase*) [108], the *Eukaryotic aspartyl protease family protein* or the *Eukaryotic aspartyl protease family protein* [109], the *TRAFlike family protein* [110]. Finally, some other genes were involved in hormonal signalization (*Auxin responsive GH3 family protein*, *Heptahelical transmembrane protein2*), in primary metabolism (*Glucose-6-phosphate dehydrogenase* playing a key role in regulating carbon flow through the pentose phosphate pathway), and in stress response (*Galactose oxidase/kelch repeat superfamily protein* [111]). Two genes encoding for transcription factors were also differentially expressed between Ti and Tf in a different way according to the plant genotype (*WRKY DNA binding protein 11* and *Basic region/leucine zipper motif 53*). The sense of expression of these genes can be correlated to the level of *P. brassicae* susceptibility of both genotypes: Yudal, known to be more resistant to clubroot than Tenor, displayed an increase of gene’s expression related to growth and disease response as potential mechanisms of resistance, whatever the microbial diversity and composition in the soil.

#### The soil microbiota composition affects the plant transcriptome

In healthy plants, the soil microbiota composition effect on plant transcriptome was similar for both genotypes: no effect at Ti and close number of DEGs at Tf. In contrast, in infected plants, only Yudal transcriptome was affected by the soil microbiota diversity, and interestingly mainly at Ti. The Yudal DEGs between L and H included a large number of genes encoding various proteins involved in plant defense, such as the CYP71A13 (phytoalexin biosynthesis), the β-glucosidase and the Nucleotide diphospho-sugar transferase (glucosinolates’ metabolism), the Pinoresinol reductase (synthesis of lignane), the oxidoreductase family protein (terpenes’ metabolism), the lectin family protein (plant defense proteins), the serine protease inhibitor and the inhibitor INAtype family protein (antimicrobial activity), the glutathione transferase F3 (transport of defense compounds), and the Lipase/Acylhydrolase superfamily protein (growth and plant defense). These proteins may represent critical early molecules in the plant defense response before disease progression.

### Complex interactions between plant/pathogen and soil microbiota

Our study aimed to decipher the interactions between plant, pathogen and the soil microbial community to better understand the mechanisms and the host/pathogen functions involved in disease modulation. We highlighted *P. brassicae* and *B. napus* DEGs between microbial environment conditions with potential functions involved in growth and pathogenicity in the pathogen, and defense in the plant. Further studies (e.g. gene inactivation) are necessary to explore if these proteins have expected functions in the Plasmodiophorids on one hand, and in *B. napus* on the other hand.

In infected plants, even the number of DEGs remained low in *B. napus*, the expression profile was pretty opposite to that of *P. brassicae* in response to soil microbiota diversity levels:

i. The plant transcriptome was more modified between H and diluted conditions for Yudal, a resistant genotype, while the pathogen transcriptome was more modified between soil microbial modalities when the host plant was Tenor, a clubroot susceptible genotype.
ii. The plant transcriptome was more modified at Ti than Tf by the soil microbial diversity, while the pathogen transcriptome was modulated later at Tf.

This host plant genotype-dependent and time-lagged response to the soil microbial composition between the plant and the pathogen transcriptomes suggest a complex regulatory scheme. The soil microbiome would modulate precociously the plant defense mechanisms in the partially resistant genotype but would have moderate or no effect in the susceptible plant, perhaps because of a too high disease level. In parallel, a direct effect of the soil microbiota composition (key-species for instance) on the pathogen could also occur in the early stages of infection, with a late visible effect on the transcriptome of the pathogen. This highlights the importance to perform studies on very early steps of infection by *P. brassicae*. Moreover, a specific microbial recruitment from the soil diversity in function of the plant genotype could also occur with subsequent consequences on pathogen metabolism in later step of its development inside the roots in interaction with endophyte microbes. These latter, differentially recruited in function of the host plant genotype, could have different effect on pathogen gene expression during its development inside the roots. In turn, the plant would affect the pathogen transcriptome by modulating or not some genes involved in growth and pathogenicity. Mutant approaches (plant and pathogen) could validate these hypotheses.

The mechanisms within the microbial functions present in soils rather than just the species need also to be studied. The difference in clubroot observed according to both plant genotypes and soil diversity could be in part explained by the concept of functional redundancy (defined as the overlapping and equivalent contribution of multiple species to a particular function) on the one hand, and the non-redundancy of rare soil microbes playing a key-role in ecosystem on the other hand [112]. Further thorough studies on microbial endophyte and rhizosphere species and functions present in both plant genotypes depending on microbial community composition are necessary to describe if some keystone microbial species/stains of specific bacteria and/or fungi could explain the clubroot phenotypes. This would require: (i) a more accurate taxonomic resolution and a more complete description (e.g. protist community) of the microbial soil compositions; (ii) a study of the functions expressed by microbial species, as described in some examples of molecular mechanisms leading to pathogen growth suppression on plant tissues found in the literature [113–116]. For this, metatranscriptomics approach to analyze the microbial functions expressed in roots are in progress to better understand the complex interaction plant / pathogen / microbial environment.

## Materials and methods

### Preparation of soils harboring different microbial diversity levels

The soil preparation to obtain different microbial diversity levels was performed as described in [25]. The soil was collected at the INRA experimental site La Gruche, Pacé, France, from the layer −10 to −30 cm. After homogenization, grinding, sieving and mixing with silica sand (2/3 soil, 1/3 sand), a part of the soil was gamma rays sterilized at 35 kGy and stabilized for 2 months. The unsterilized soil (100 g of dry soil) was suspended in 1 L of deionized water and used for serial dilution: undiluted (10^0^, High diversity level [H], considered as the reference), diluted at 10^−3^ (Medium diversity level [M]) or 10^−6^ (Low diversity level [L]). Three dilution processes were performed corresponding to 3 biological replicates. The sterilized soil (2.5 kg per bag) was inoculated with 300 mL of each dilution (H, M, L) and incubated in the dark at 18°C and 50% humidity for 49 days. Every week, microbial respiration and recolonization were facilitated when opening the bags under hood. The recolonization was followed by a microbiological count of formed cultivable colonies during the incubation period (S1 Fig).

### Molecular characterization of soil bacterial and fungal communities

After recolonization and before sowing, the three microbial modalities were analyzed for their physicochemical composition at the Arras soil analysis laboratory (LAS, INRA, Arras, France) (S1 Table) and for their microbial diversity. The GnS-GII protocol was used for extraction of DNA from soil samples [117]. Briefly, DNA was extracted from 2 g of dry soil, and then purified by PVPP column and Geneclean [28]. PCR amplification and sequencing were performed at the GenoScreen (Lille, France) using the Illumina MiSeq ‘paired-end’ 2×250 bases (16S) for bacteria and Illumina MiSeq ‘paired-end’ 2×300 bases (18S) for fungi as described previously [25, 28]. The protist diversity was not included in the analysis. After read assembly, sequences were processed with the GnS-PIPE bioinformatics developed by Genosol platform [118, 119]. By performing high-quality sequence clustering, Operational Taxonomic Units (OTUs) were retrieved and taxonomic assignments were performed comparing OTUs representative sequences against dedicated reference databases from SILVA [120]. The cleaned data set is available on the European Nucleotide Archive database system under the project accession number PRJEB36457. Soil samples accession numbers range from ERR3842608 to ERR3842625 for 16S and 18S rDNA.

The alpha diversity of the communities was analyzed. To compare bacterial or fungal composition among three soil preparations, the richness of these communities was characterized by the number of OTUs found in each soil. As metric of taxonomy diversity, the Shannon diversity index was also determined (package ‘vegan’ [121]). Since values were conformed to normality assumptions, linear models LMM function ‘lmer’, package ‘lme4’ [122]) were used to examine differences between soil preparation for these measures. When needed, pairwise comparisons of least squares means (package ‘lsmeans’ [123]) and a false discovery rate correction of 0.05 for P-values [124] were performed.

In order to analyse the bacterial and fungal community structure (beta diversity), principal coordinate analysis (PCoA) was performed on a Bray-Curtis dissimilarity matrix, obtained from OTUs data, which were normalized using a 1‰ threshold and log2-transformed (package ‘vegan’ [121]). A type II permutation test was performed on the PCoA coordinates to compare the community structure of the H, M and L soils (package ‘RVAideMemoire’ [125]).

### Plant material and pathogen inoculation

The oilseed rape genotypes Tenor and Yudal and the eH isolate of *P. brassicae* belonging to pathotype P1 [39, 126, 127] were used in this study. Yudal and Tenor genotypes were chosen because previous assay in our lab showed they display different responses to clubroot infection: Tenor was more susceptible than Yudal to eH. Both *B. napus* genotypes were grown in each of the three soils (harboring H, M or L microbial diversities). For this, seeds of oilseed rape were sown in pots filled with 400 g of experimental soils. Pots were placed in a climatic chamber, in a randomized block design with the three modalities (H, M, L) and three replicates by dilution factor. For each oilseed rape genotype, eight plants per soil microbial modality and per replicate were used. Plants were either not inoculated (healthy plants) or inoculated with a resting spore suspension of the *P. brassicae* eH isolate. For inoculum production, clubs propagated on the universal susceptible host Chinese cabbage (*B. rapa* ssp *pekinensis* cv. Granaat) were collected, homogenized in a blender with sterile water and separated by filtration through layers of cheesecloth. The resting spores were then separated by filtration through 500, 100 and 55 µM sieves to remove plant cell debris. The spore concentration was determined with a Malassez cell and adjusted to 1.10^7^ spores.mL^−1^. Plant inoculation was done as described in [128]: seven-day-old seedlings were inoculated by pipetting 1 mL of the spore suspension at 1.10^7^ spores.mL^−1^ to the bottom of the stem of each seedling. The plants were maintained at 22°C (day) and 19°C (night) with a 16h photoperiod, and watered periodically from the top with a Hoagland nutritive solution to provide nutrients and to maintain a water retention capacity of 70 to 100%.

### Phenotyping: plant characterization and disease assessment

Roots and aerial parts were sampled at two times: 28 days after inoculation (dai) (intermediary time, Ti) for both genotypes, and 36 dai and 48 dai for Tenor and Yudal (final time, Tf), respectively. The final time was chosen to have clearly visible galls on the primary and lateral roots.

At each sampling date and for each replicate, the aerial parts of 8 plants were cut, dried and weighted. As one of the three infected replicates at the final time for Tenor in L soil displayed no clubroot symptoms in any of the 8 plants, indicating that the inoculation of these plants was not successful, this sample was removed for all the analyses. The roots were cut below the collar (in the soil depth from −1 to −6 cm), separated from soil, and washed twice in sterile water by vortexing 10 sec. Then the roots were transferred in a petri dish, cut into small pieces, and frozen in liquid nitrogen then stored at −80°C. After lyophilization, the dry root biomass was measured and the powder was kept until nucleic acid extraction (DNA for pathogen quantification and RNA for RNAseq analyses).

Disease was assessed at each sampling date after inoculation with *P. brassicae*. First, clubroot symptoms were evaluated by a disease index calculated with the scale previously described by Manzanares-Dauleux et al. [128]. Secondly, 1 µL of DNA extracted from root samples (see 2.5.) was used for quantitative PCR on the LightCycler® 480 Real-Time PCR System (Roche) to quantify *P. brassicae* amount. For this, a portion (164 bp) of the target 18S gene was amplified with the following primers: 5’-ttgggtaatttgcgcgcctg-3’ (forward) and 5’-cagcggcaggtcattcaaca-3’ (reverse). Each reaction was performed in 20 µL qPCR reaction with 10 µL of SYBR Green Master Mix (Roche), 0.08 µL of each primer (100 µM) and 1 µL of total DNA as template. The PCR conditions consisted of an initial denaturation at 95°C for 5 min, followed by 45 cycles at 95°C for 10 s and 64°C for 40 s. Standard curves were constructed using serial dilutions of *P. brassicae* DNA extracted from resting spores. Quantitative results were then expressed and normalized as the part of the *P. brassicae* mean DNA content in the total root-extracted DNA.

To compare the aerial and root biomasses between modalities, linear models were used (LMM function ‘lmer’, package ‘lme4’ [122]). A Wald test (α = 5%) was applied for evaluating the soil effect in the LMM model. Least Square Means (LSMeans) were calculated using the ‘lsmeans’ function of the ‘lsmeans’ package [123], and the false discovery rate correction for P-values [124]. Pairwise comparisons of LSMeans were performed with the Tukey test (α = 5%), using the ‘cld’ function of the ‘lsmeans’ package.

Disease data were analyzed using a likelihood ratio test on a cumulative link model (CLMM; ‘clmm’ function, ‘ordinal’ package). LSMeans and pairwise comparisons of LSMeans were performed as described for biomasses’ analyses.

### Nucleic acids isolation from roots

At each time-point, the lyophilized roots from the 8 pooled plants of each genotype and each treatment (with and without *P. brassicae*) were used for nucleic acid extraction. DNA was extracted from 30 mg of lyophilized powder root samples with the NucleoSpin Plant II Kit (Masherey-Nagel) following the manufacturer’s instructions. After verification of the DNA quality on agarose gel and estimation of the quantity with a Nanodrop 2000 (Thermoscientific), it was used for *P. brassicae* quantification.

Total RNA was extracted from 20 mg of lyophilized powder with the Trizol protocol (Invitrogen). RNA purity and quality were assessed with a Bioanalyser 2100 (Agilent) and quantified with a Nanodrop (Agilent).

### Library construction and Illumina sequencing

RNA-seq analysis was performed on RNA extracted from roots tissues of two *B. napus* genotypes infected or not with resting spores of *P. brassicae* (eH isolate) grown in the three different soils (H, M, L), for three biological replicates, at Ti and Tf.

The TruSeq Stranded mRNA Library Sample Prep Kit (Illumina) was used for library construction. Library pair-end sequencing was conducted on an Illumina HiSeq4000 (Genoscreen, Lille, France) using 2×150 bp and resulting in 2861 paired-end millions of reads. Briefly, the purified mRNA was fragmented and converted into double-stranded cDNA withy random priming. Following end-repair, indexed adapters were ligated. The cDNA fragments of ∼350 pb were purified with AMPure beads XP and amplified by PCR to obtain the libraries for sequencing. The libraries were multiplexed (six libraries per lane) and sequenced. The cleaned data set is available on the European Nucleotide Archive database system under the project accession number PRJEB36458. Samples accession numbers range from ERR3850126 to ERR3850197.

### Mapping of sequenced reads, assessment of gene expression and identification of differentially expressed genes

The read quality was undertaken for the quality scores of Q28 and for the read length of 50 nucleotides using PrinSeq. In order to use a combined host-pathogen genome as reference for alignment, the genomes of eH *P. brassicae* [39] and *B. napus* [129] were concatenated, as well as the corresponding annotation files. The high-quality reads were aligned to the concatenated files using STAR 2.5.2a_modified. Non-default parameters were minimum intron length 10, maximum intron length 50 000 and mean distance between paired ends-reads 50 000. For the reads which can align to multiple locations (parameters set for a maximum of 6 locations), a fraction count for multi mapping reads was generated. Thanks to genome annotation files, the mapped sequencing reads were assigned to genomic features using featureCounts v1.5.0-p1, and counted. After filtering of the read counts below the threshold value (at least 0.5 counts per million in 3 samples), the count reads were then normalized with the Trimmed Mean of M values (TMM method). Concerning the *P. brassicae* reads, as the number of reads in the libraries at Ti was much smaller than at the final time (due to the differences in the infection rate and progression of the pathogen between the sampling times), the normalization was performed for Ti separately from Tf. So, analyses of *P. brassicae* were specific of each sampling time, preventing the data comparison between the time-points. On the contrary, for *B. napus* reads, the normalization was performed on total libraries, allowing a kinetic analysis of plant transcriptome.

Differential expression analysis was performed using the EdgeR package in R. The Differentially Expressed Genes (DEGs) with FDR ≤ 0.05 from specific comparison lists were selected for analysis. The functional annotation of DEGs was performed with Blast2GO 4.1.9 software. Heat maps were generated using the ‘heatmap3’ package and Venn Diagrams using the ‘VennDiagram’ packages in R.

## Supporting information captions

S1 Fig. Microbiological follow up based on the Colony Forming Units (CFU) method during the incubation period for bacteria (A) and fungi (B). H, High diversity modality; M, Medium diversity modality; L, Low diversity modality.

S2 Fig. Description of the main bacterial and fungal composition in the three soils. Average relative abundance (RA ± SEM) of the most abundant bacterial phyla (A), genera (B), OTUs (C), and fungal phyla (D), genera (E), OTUs (F) are shown in High (H), Medium (M) and Low (L) soil microbial diversities. For each soil, the number of replicates is n=3.

S3 Fig. Overview of all *P. brassicae* transcriptome samples. A. Heatmaps of *P. brassicae* gene expression based on normalized data of expression values. The heatmaps are based on total reads counts for *P. brassicae* at Ti and Tf for the 3 microbial soil diversities (H, High; M, Medium, L, Low), the two plant genotypes (T, Tenor; Y, Yudal) and correspond to the mean of the three replicates. B. Hierarchical Cluster Analysis (HCA) of the filtered and normalized counts in the dual-RNAseq analysis. The analyses are shown for *P. brassicae* reads at Ti and Tf for the 3 soil microbial diversities (H, High; M, Medium; L, Low), the two plant genotypes (T, Tenor; Y, Yudal), and the three replicates (a, b, c).

S4 Fig. Overview of all *B. napus* transcriptome samples. Hierarchical Cluster Analysis (HCA) of the filtered and normalized counts in the dual-RNAseq analysis in healthy plants (A) and infected plants (B). The analyses are shown for *B. napus* reads at Ti and Tf, for the 3 soil microbial diversities (H, High; M, Medium; L, Low), the two plant genotypes (T, Tenor; Y, Yudal), and the three replicates (a, b, c).

S5 Fig. Number of *B. napus d*ifferentially expressed genes (DEGs) in function of the host plant genotype for each soil microbial diversity level when not infected (A) or infected by *P. brassicae* (B). The Venn diagram shows the number of significantly DEGs (P < 0.05) according to the host *B. napus* genotypes (T, Tenor; Y, Yudal) infected or not, for each soil microbial diversity level (H, High; M, Medium; L, Low) at the sampling dates Ti and Tf.

S6 Fig. Number of *B. napus* differentially expressed genes (DEGs) in function of the interaction stage for each soil microbial diversity level. The Venn diagrams show the total number of significantly DEGs (P < 0.05) in the *B. napus* genotypes (T, Tenor; Y, Yudal), healthy (A) or infected by *P. brassicae* (B), at each soil microbial diversity level (H, High; M, Medium; L, Low), between Ti and Tf.

S7 Fig. Differentially expressed genes (DEGs) in both infected *B. napus* genotypes according to the infection’s stage whatever the soil microbial diversity. A. The Venn diagram shows the number of significantly DEGs (P < 0.05) common in both *B. napus* genotypes (T, Tenor; Y, Yudal), and common in the three soil microbial diversity levels (H, High; M, Medium; L, Low), which are down (<) or up (>) regulated at Ti compared to Tf. B. Heatmaps of the 40 genes surrounded by a grey circles in the figure A. The expression is based on normalized data of expression values (T, Tenor; Y, Yudal; H, M, L, High, Medium, Low soil microbial diversity levels).

S1 Table. Main physicochemical characteristics of the three soils used in this study.

S2 Table. Description of the *P. brassicae* genes differentially expressed between H and M at Tf when infecting Yudal (−1: genes underexpressed at H compared to M; 1: genes overexpressed at H compared to M).

S3 Table. Description of the *P. brassicae* genes differentially expressed between the different soil microbiota diversity levels at Tf when infecting Tenor. A. Description of the *P. brassicae* genes differentially expressed between H and M at Tf when infecting Tenor (−1: genes underexpressed at H compared to M; 1: genes overexpressed at H compared to M). B. Description of the *P. brassicae* genes differentially expressed between H and L at Tf when infecting Tenor (−1: genes underexpressed at H compared to L; 1: genes overexpressed at H compared to L). C. Description of the *P. brassicae* genes differentially expressed between H and M and between H and L at Tf when infecting Tenor (−1: genes underexpressed at H compared to M or L; 1: genes overexpressed at H compared to M or L).

S4 Table. Description of the P. brassicae genes differentially expressed between H and M at Tf in an opposite sense when infecting Yudal or Tenor (−1: genes underexpressed at H compared to M; 1: genes overexpressed at H compared to M).

S5 Table. Effect of soil microbiota diversity levels on infected Yudal gene expression. A. Description of the 64 B. napus Yudal genes differentially expressed between H and L at Ti when infected by P. brassicae (−1: genes underexpressed at H compared to L; 1: genes overexpressed at H compared to L). B. Description of the 23 B. napus Yudal genes differentially expressed between H and L at Tf when infected by P. brassicae (−1: genes underexpressed at H compared to L; 1: genes overexpressed at H compared to L).

## Acknowledgments

We thank the Biological Resources Center *BrACySol* (INRA Rennes, France) for providing the Brassica seeds. This work was supported by grants from the Plant Health and Environment division and the Plant Biology and Breeding division of the French National Research Institute for Agriculture, Food and the Environment (INRAE).

## Author Contributions

**Conceptualization**: Stéphanie Daval, Christophe Mougel

**Data curation**: Kévin Gazengel, Arnaud Belcour

**Formal analysis**: Stéphanie Daval, Kévin Gazengel, Arnaud Belcour, Lionel Lebreton

**Funding acquisition**: Stéphanie Daval, Alain Sarniguet

**Investigation**: Stéphanie Daval, Kévin Gazengel, Juliette Linglin, Anne-Yvonne Guillerm-Erckelboudt, Lionel Lebreton, Christophe Mougel

**Methodology**: Stéphanie Daval, Kévin Gazengel, Arnaud Belcour, Juliette Linglin, Anne-Yvonne Guillerm-Erckelboudt, Lionel Lebreton, Christophe Mougel

**Project administration**: Stéphanie Daval

**Resources**: Stéphanie Daval, Kévin Gazengel, Lionel Lebreton, Christophe Mougel

**Supervision**: Stéphanie Daval, Christophe Mougel

**Validation**: Stéphanie Daval, Maria J. Manzanares-Dauleux, Christophe Mougel

**Visualization**: Stéphanie Daval, Kévin Gazengel, Lionel Lebreton

**Writing – original draft**: Stéphanie Daval

**Writing – review and editing**: Stéphanie Daval, Maria J. Manzanares-Dauleux, Lionel Lebreton, Christophe Mougel

